# Toll-like receptor-1 (TLR-1) activation reduces immunoglobulin free light chain production by multiple myeloma cells in the context of bone marrow stromal cells and fibronectin

**DOI:** 10.1101/2024.09.02.610867

**Authors:** Jahan Abdi, Frank Redegeld

## Abstract

Studies over the past years have provided evidence that Toll-like receptor (TLRs) activation in multiple myeloma (MM) cells induces heterogeneous functional responses including cell growth and proliferation, survival or apoptosis. These effects have been suggested to be partly due to increase in secretion of cytokines such as IL-6 or IFNα among others from MM cells following TLR activation. However, whether triggering of these receptors also modulates production of immunoglobulin free light chains (FLCs) in MM cells has never been investigated. FLCs contribute largely to MM pathology. Here we explored the effect of TLR1 ligand (Pam3CSK4) alone or combined with bortezomib (BTZ) on production of FLCs in human myeloma cell lines, L363, OPM-2, U266 and NCI-H929 in the absence or presence of bone marrow stromal cells (BMSCs) or fibronectin (FN) to examine the influence of bone marrow microenvironment. Adhesion to BMSCs or FN increased secretion of FLC in MM cells. Pam3CSK4 decreased FLC production in the presence or absence of BMSCs or FN and this effect was enhanced in combination with BTZ. However, the level of reduction was lower in the presence of BMSCs or FN. Our findings imply that activation of TLR1 downregulates FLC production in MM cells even in the context of bone marrow microenvironment components and suggest that some TLRs such as TLR1 might be considered a therapeutic target especially in combined treatment protocols in MM.

## Introduction

Within the bone marrow microenvironment, MM cells closely interact with BMSCs or extra cellular matrix proteins such as FN, collagen, laminin, and tenascin [1, 2]. This interaction plays a key role in maintaining the survival and drug resistance of MM malignant clones, mediated mostly by direct contact (adhesion) or secretion of a plethora of cytokines and growth factors by BMSCs or MM cells, including IL-6, VGEF, IGF-1, HGF, IL-3 and RANK [3, 4]. These cytokines are also known to contribute to angiogenesis and osteolytic bone lesions in MM [3, 4].

MM cells are dependent on microenvironmental cues to survive [5, 6], however, in advanced stages of the disease tumor cells may gain multiple gene mutations which could render them independent of stromal components [7-9]. As potential extracellular signals, Toll-like receptor (TLR) ligands were suggested to play important modulatory roles in growth, proliferation and viability of MM cells which display high but heterogeneous expression pattern of these receptors [10-15]. It is conceivable that TLR activation in MM bone marrow microenvironment could also influence MM cells interaction with BMSCs or FN leading to modulation of MM cells secretory functions. Of note, MM malignant clones produce immunoglobulin free light chains (FLCs) which are known to contribute significantly to MM complications [16, 17]. However, whether TLR triggering in MM bone marrow microenvironment modulates FLC secretion by MM cells is not known.

The biologic pattern of FLCs production has not been convincingly explored in MM experimental models following drug treatment. In around 20% of MM patients referred to as light chain type myeloma, only one monoclonal immunoglobulin light chain is detectable, whilst in most types of MM excess light chain proteinuria might occur. When re-absorption of FLCs exceeds the maximum potential of renal tubular system, they form in situ protein complexes resulting in nephropathy [18]. Therefore, targeting mechanisms controlling the production of FLCs by the malignant clone would be an efficient therapeutic approach.

We have previously shown that treatment of human myeloma cell lines and MM primary cells with Pam3CSK4 (TLR1/2 ligand) sensitized them to cytotoxic effect of BTZ in the context of BMSCs or FN [19, 20]. In the present research we sought to explore the effect of Pam3CSK4 (alone or combined with BTZ) on FLC production by HMCLs interacting with FN or BMSCs. We showed that TLR1 activation by Pam3CSK4 decreased production of FLCs in HMCLs, while the effect of Pam3CSK4 was slightly attenuated following adhesion of HMCLs to FN or BMSCs. Further studies are required to uncover the molecular mechanisms underlying TLR-mediated modulation of FLC in MM cells.

## Materials & methods

### Cell lines and cell culture

Human multiple myeloma cell lines (HMCLs), L363, OPM-2, U266, and NCI-H929, were provided by Hematology department of Utrecht University Medical Center [21, 22], human bone marrow stromal cell line, HS.5, was obtained from American Type Culture Collection (ATCC). HMCLs were maintained in RPMI medium supplemented with 5-10 % FBS, 2mM glycine, and intermittently with antibiotics. HS-5 was maintained in DMEM medium supplemented with 10% FBS, and intermittently with antibiotics.

### Chemicals

TLR1/2 ligand (Pam3CSK4) was obtained from Invivogen and dissolved in sterile water according to manufacturer’s instructions to make a 1 mg/ml stock. Bortezomib (BTZ) was from LC Laboratories and was dissolved in DMSO as a 100mM stock. The final DMSO concentration never exceeded 0.01% in all experimental conditions. Human plasma-derived fibronectin was from Sigma.

### Cell stimulation

For cells stimulation we followed a sequential treatment procedure: First, HMCLs were harvested from cultures and incubated with a range of Pam3CSK4 doses (0.05-2.5µg/ml) for 24h. Then, cells were washed and re-suspended in fresh RPMI medium and seeded (or not) onto fibronectin- or BMSCs-coated wells of a 96-well plate. In other experiments, wells were exposed to specific dose of BTZ as explained below.

### Co-culture of MM cells with HS-5

To assess FLC production by TLR1-activated HMCLs in the context of bone marrow stromal cells, 100,000 HS-5 cells were seeded on 12-well plates to achieve 70% confluency in almost 40h. Five hundred thousand cells from HMCLs pre-activated for 24 h with Pam3CSK4 were washed, re-suspended in fresh serum-free RPMI medium and exposed to stromal cell-coated wells for 2 h. Unattached cells were removed and fresh medium containing protein was added and plates were further incubated for 24 h. At the end of incubation, supernatants were collected and assayed for FLC concentrations. In separate experiments, to assess the combined effect of BTZ+Pam3CSK4 on FLC production in HS-5 context, before seeding the HMCLs, they were incubated with 1µM of BTZ in protein-containing medium for one h (*acute exposure*), washed and resuspended in warm drug-free medium, added to stromal layer and incubation was further extended to 24 h.

### Culture of HMCLs on FN-coated microwells

To analyze the effect of Pam3CSK4 on FLC production by HMCLs in the context of FN, 12-well plates were coated with 20µg/ml FN overnight at 4°C. Plates were blocked with sterile heat-denatured BSA (10mg/ml in PBS) for 1 h at room temperature and washed. HMCLs pre-activated with Pam3CSK4 as described above were washed, re-suspended in serum-free RPMI and seeded on coated plates for 1 h. Unattached cells were removed, and fresh medium containing protein was added, and the plates were further incubated for 24 h. In separate experiments, to assess the combined effect of BTZ+Pam3CSK4 on FLC production in FN context, BTZ (5nM) in RPMI medium (containing protein) was added after removing the unattached cells and incubation extended to 24 h (*chronic exposure*).

### ELISA

Kappa or lambda FLCs were assayed as previously described [23]. Briefly, 96 well plates were coated overnight at 4°C with goat anti-mouse IgG antibody in bicarbonate buffer. Subsequently, plates were blocked for 1 h (RT) and incubated with mouse-anti human kappa or lambda Ig-FLC mAbs (obtained from Dr. A. Solomon, Tennessee). After incubation with different dilutions of samples and standards (The Binding Site), plates were incubated with HRP-labelled goat F(ab’)2 anti-human kappa or lambda Ig light chain Abs (AHI1804 and AHI1904, respectively, Biosource, USA). Finally, the reactions were developed using TMB and measured through an ELISA plate reader (BioRad).

### Statistics

We used 2-way ANOVA in GraphPad software for data analysis and *p*<0.05 was considered as significant.

## Results

### Baseline production of FLC by HMCLs

We first determined the baseline concentration of FLC in all the cell lines and found heterogenous pattern of LC isotypes. As shown in table 1, L363, OPM2, U266, were lambda (λ) light chain producers, and NCI-H929 was a kappa (κ) light chain producer.

**Table 1.**
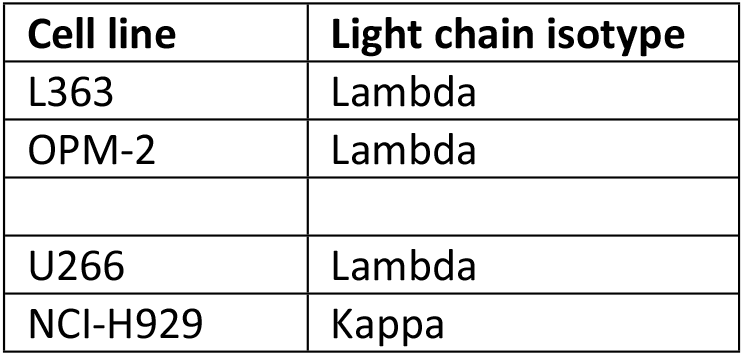
Production of different light chain isotypes by HMCLs at baseline.

### TLR1 activation in HMCLs reduces FLC secretion

L363, OPM-2, U266 (lambda light chain producers) and NCI-H929 (kappa light chain producer) HCMLs (Table 1) were analyzed for FLC production following Pam3CSK4 treatment. HMCLs were stimulated with a range of Pam3CSK4 concentrations (0.05-5.0µg/ml) for 24h and FLC concentration was measured in culture supernatants. Our previous study showed that Pam3CSK4 would induce apoptosis in HMCLs at 2.5µg/ml or higher [20]. Therefore, we investigated if the inhibitory effect of Pam3CSK4 on FLC production was due to increased cell death. To this aim, for L363 and OPM2 cells we determined viability of cells related to each Pam3CSK4 concentration using propidium iodide staining followed by flow cytometry. Depicted in figure 1, all Pam3CSK4 doses reduced FLC production in both cell lines; however, it generated a bell-shaped pattern. While further investigation is required to elucidate this dynamic nature of TLR-1 activation effect on FLC production in MM cells, it could be safe to assume that at lower end of the range (0.05-0.25µg/ml) Pam3CSK4 specifically reduces FLC production, but at higher doses FLC reduction is mostly due to cytotoxicity. As can be seen in figure 1, Pam3CSK4 significantly decreased FLC production at the lowest dose (0.05µg/ml) while the viability is remarkably high, and the same dose in separate experiments also significantly reduced FLC level in the presence or absence of FN or BMSC, although the response was not significant in U266 (**Figs 2,3**).

**Figure 1.**
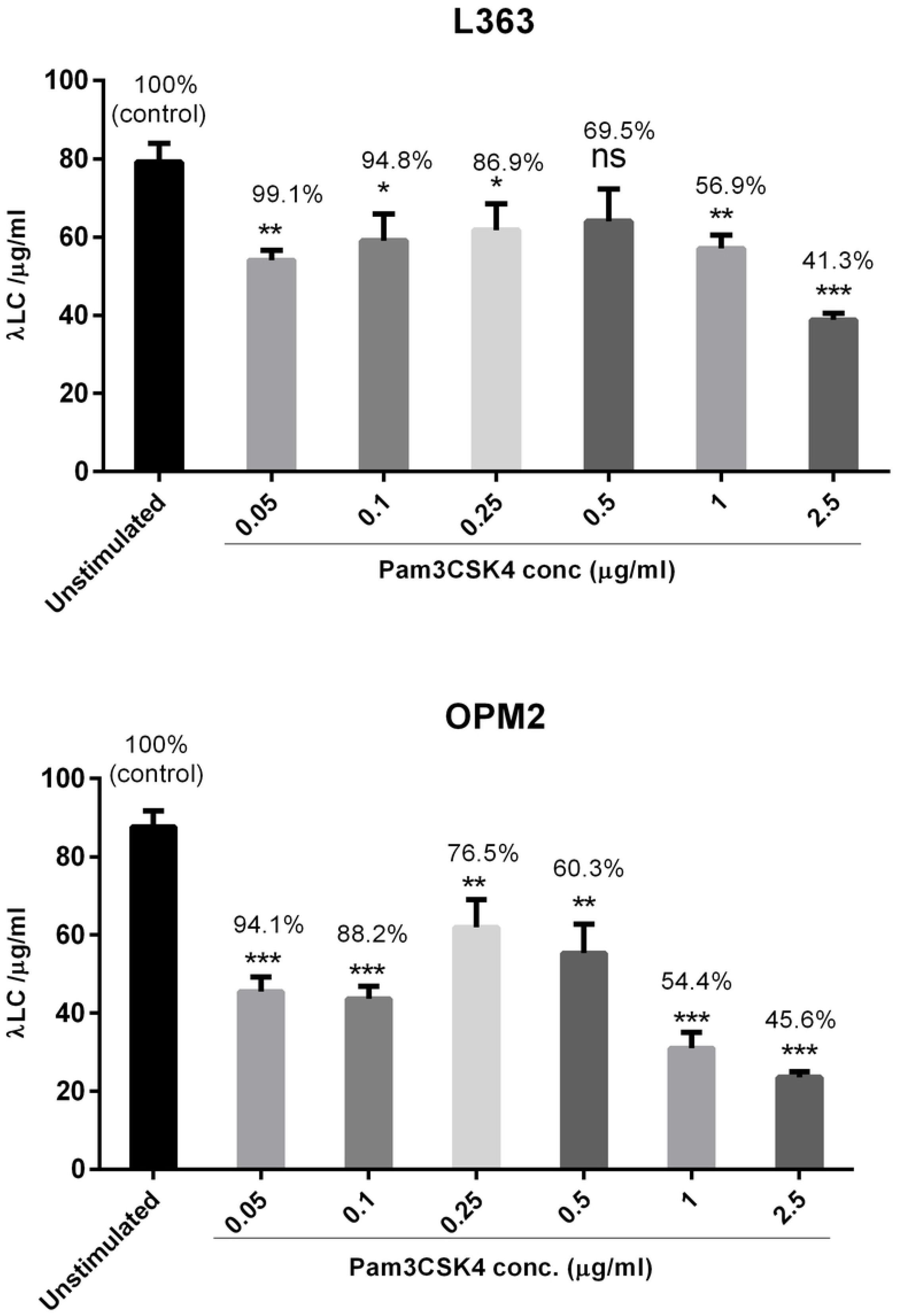
Effect of Pam3CSK4-induced cell death on FLC production by HMCLs. L363 and OPM2 cells were stimulated with a wide range of Pam3CSK4 concentrations (0.05-2.5µg/ml) for 24 h. Supernatants were collected for FLC analysis, and cells were applied to PI FACS staining to determine % viability of cells and whether FLC production was affected by cell death (% on top of the bars indicate viability of cells in each condition)

**Figure 2.**
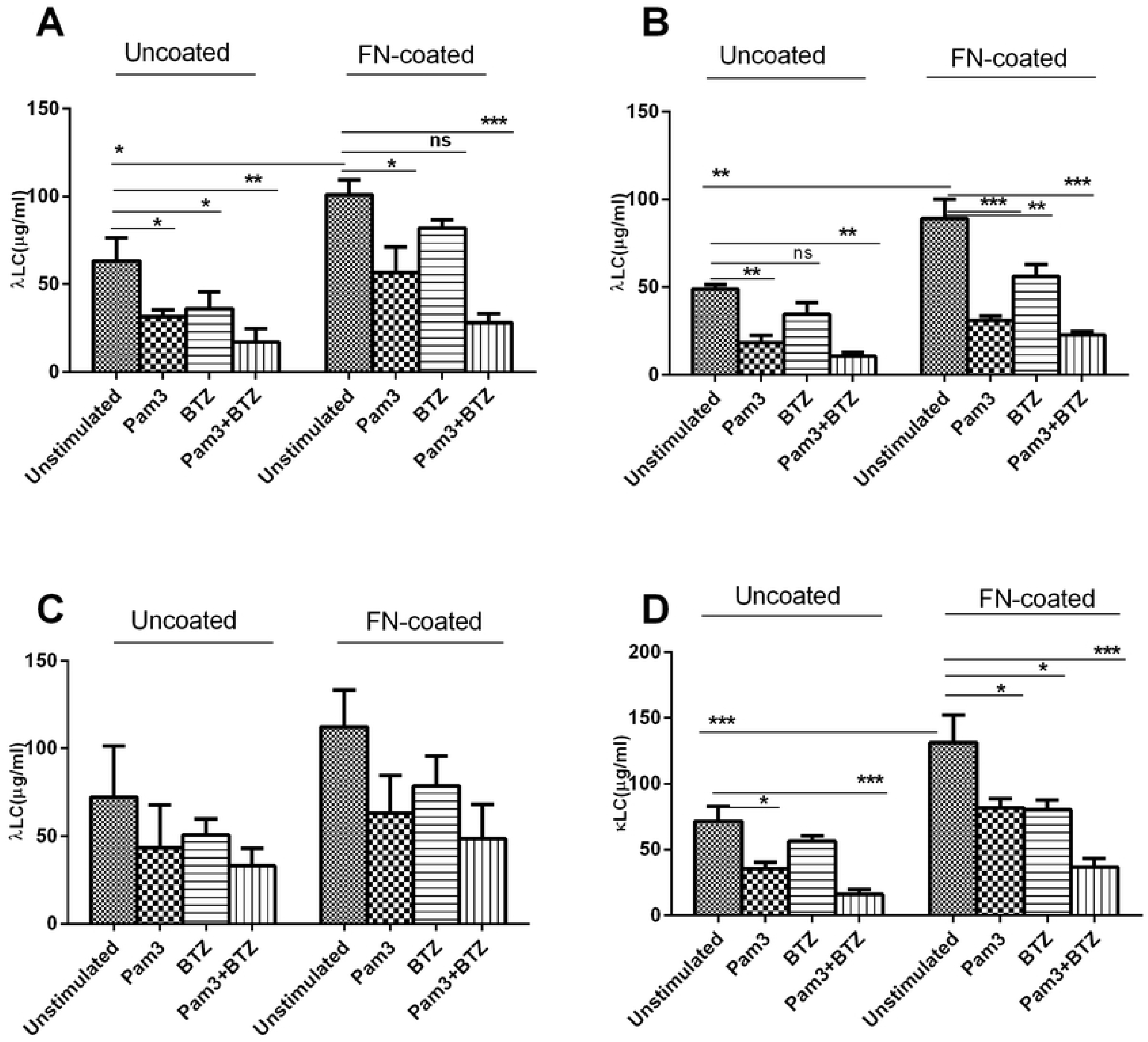
The effect of Pam3CSK4 on FLC production in HMCLs : λ in L363 (**A**), OPM-2 (**B**), U266 (**C**), and κ in NCI-H929 (**D**) after exposure to BTZ in the context of FN. HMCLs were stimulated (or not) with 0.05µg/ml Pam3CSK4 for 24 hours, washed, adhered (or not) to FN and exposed (or not) to 5nM BTZ for 24h as explained in materials and methods. Data are the mean±SD from analysis of three separate experiments. **\*** *p < 0*.*05*, **\*\*** *p <0*.*01*, **\*\*\*** *p <0*.*001*.

**Figure 3.**
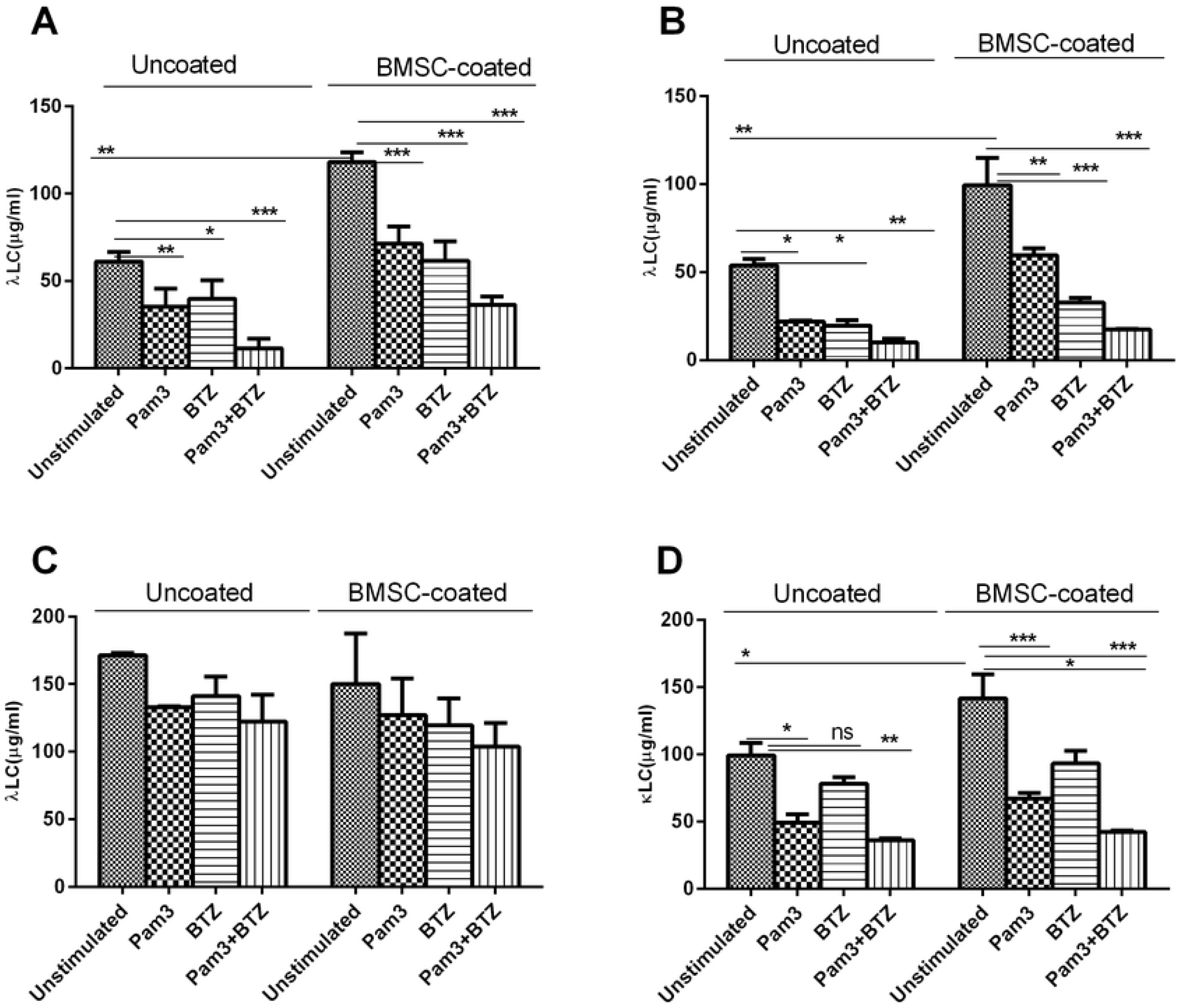
The effect of Pam3CSK4 on FLC production in HMCLs: λ in L363 (**A**), OPM-2 (**B**), U266 (**C**), and κ in NCI-H929 (**D**) after exposure to BTZ in the context of BMSCs. HMCLs were stimulated (or not) with 0.05µg/ml Pam3CSK4 for 24 h, washed, adhered (or not) to BMSCs and exposed (or not) to 1µM BTZ for one h (more explanation in the text). Data are the mean±SD from analysis of three separate experiments. **\*** *p < 0*.*05*, **\*\*** *p <0*.*01*, **\*\*\*** *p <0*.*001*.

### Adhesion to BMSCs or FN increases secretion of FLCs in HMCLs

It has been shown that adhesion of MM cells to extracellular matrix proteins such as FN or to BMSCs induces their proliferation and renders them resistant to drugs, which was originally termed cell adhesion-mediated drug resistance (CAMDR) [24, 25]. To show that such adhesion also modulates secretion of FLC by MM cells, HMCLs were co-cultured with HS.5 cells or cultured on FN-coated microwells for 24 h. As shown in figures 2,3, MM cells increased their production of FLCs following adhesion to FN or HS.5 cells (uncoated, unstimulated vs coated, unstimulated). In U266 cell line, results did not reach significance probably due to a higher level of variation between experiments. These observations imply that BMSCs might play a regulatory role in FLC secretion by MM cells, however, further mechanistic studies are required to support this.

### Pam3CSK4-induced reduction in FLCs is enhanced in combination with BTZ in the presence or absence of bone marrow microenvironment

It is well established that adhesion of MM cells to bone marrow stroma components such as FN and BMSCs induces secretion of various cytokines in MM cells and renders them resistant to drugs [3, 25]. Having shown that the above adhesion upregulates FLC production in HMCLs, we sought to explore whether Pam3CSK4 alone or in combination with BTZ would affect FLC production by HMCLs while these cells are adhered to FN or BMSCs. Again, as shown in figures 2 and 3, BTZ significantly decreased FLC production in L363, OPM2 and NCI-H929 cell lines, in the presence or absence of FN or BMSCs, but this inhibitory effect of BTZ was not significant in U266 cells. Also, by comparing the two conditions, uncoated vs coated, in all cell lines it can be seen that inhibitory effect of BTZ or Pam3CSK4 is slightly attenuated in adhered cells due to the impact of BMSCs or FN. Combined treatment with Pam3CSK4+BTZ showed at least an additive effect in reducing FLC production although in adhered cells this inhibitory effect was also lower. Furthermore, to see if FLC reduction by BTZ is not due to cell death, in a separate experiment 10000 cells each of L363 and OPM2 cell lines were exposed to a dose range of BTZ (0.25-5.0nM) for 24h in a 96-well plate. At the end of incubation, the number of live cells for each dose was determined in parallel with FLC measurement (supp. Fig1). The results imply that FLC reduction is in correlation with live cell number, as BTZ dose increases, FLC is decreased and so does the number of viable cells. However, the effects of cell death on FLC production especially at BTZ doses above 2.0nM cannot be excluded. Taken together, our findings indicate that Pam3CSK4 (TLR1 activation) in HMCLs diminishes FLC production in the presence or absence of bone marrow microenvironment components and this effect would be enhanced in combination with BTZ at least in an additive fashion.

## Discussion

This study is the first to report that TLR activation modulates FLC production in myeloma cell lines, L363, OPM-2, U266 and NCI-H929. It showed that Pam3CSK4 significantly decreased FLC production by all HMCLs. In our previous work, we demonstrated that Pam3CSK4 at 2.5µg/ml induced cell death in MM cells [20]. Here we analyzed cell death by propidium iodide uptake using FACS staining of L363 and OPM2 cells treated with a range of Pam3CSK4 concentrations and in parallel FLC concentration was measured. Pam3CSK4 significantly reduced FLC production at the lowest dose (0.05µg/ml) with no effect on cell viability. This suggests the inhibitory effect of Pam3CSK4 on FLC production in HMCLs is more prominent.

Numerous studies have demonstrated the protective role of BMSCs and FN on MM cells leading to cell adhesion mediated drug resistance (CAMDR) phenotype [3, 26-31]. However, no study has yet explored whether the above components also influence the pattern of FLC production in MM cells. Here, we showed that adhesion of MM cells to FN or BMSCs significantly upregulated FLC secretion by MM cells suggesting that adhesion-mediated signaling is probably involved in FLC regulation in MM microenvironment. Furthermore, Pam3CSK4 effectively reduced FLC level in the presence or absence of bone marrow microenvironment components and this effect was significantly enhanced in combination with BTZ at least additively. On the other hand, the extent of FLC reduction in adhered cells (FN and BMSCs) was lower compared to non-adhered cells implying that FLC inhibitory effects of BTZ or BTZ+PamCSK4 could be hampered due to increased secretion of FLC by MM cells in the presence of bone marrow stroma. components of bone marrow microenvironment may hamper BTZ or Pam3CSK4 were not adhered to above components. Our descriptive study provides some new insights into the impact of TLR activation on FLC production by MM cells, pattern of FLC production by MM cells in the presence of bone marrow microenvironment components, and therapeutic potential of Pam3CSK4+BTZ in terms of FLC reduction. Notably, development of resistance to bortezomib in MM patients has been reported [32-34]. It remains to be explored mechanistically how TLR activation impacts FLC production, and whether/how integrin activation (cell-cell adhesion) could regulate FLC production, understanding these mechanisms will help identifying new therapeutic targets in MM microenvironment. Finally, our study suggests further investigation into pre-clinical application of Pam3CSK4 alone or in combination with BTZ in MM *in vivo* studies.

## Author Contributions

JA designed the study, performed experiments, did data collection and analysis, and wrote the paper. FR contributed to design of the study and data analysis, supervised the project and reviewed/edited the paper.

## Funding Sources

This study was not supported by any specific sponsor or funder.

**Supplementary figure 1. Effect of BTZ-induced cell death on FLC production in MM cells**. 10000 cells each of L363 and OPM2 cell lines were exposed to a dose range of BTZ (0.25-5.0nM) for 24h in a 96-well plate. At the end of incubation, the number of live cells for each dose was determined in parallel with FLC measurement.

